# Hippocampome.org, a resource for subicular neuron types and beyond

**DOI:** 10.64898/2025.12.30.697062

**Authors:** Carolina Tecuatl, Giorgio A. Ascoli

## Abstract

To establish the relationship between circuit organization and information processing, many neuroscientists find it useful to reason in terms of neuron types. Hippocampome.org uses axonal and dendritic morphology as a foundational approach to classify neurons in the rodent hippocampal formation, including dentate gyrus, Cornu Ammonis, subiculum, and entorhinal cortex. For each identified neuron type, this open access knowledge base annotates essential properties, such as main neurotransmitter, membrane biophysics, firing patterns, molecular expression, and cell counts. Moreover, Hippocampome.org quantifies circuitry in terms of directional connection probabilities and synaptic signals between interacting neuron types. All properties are directly linked to peer reviewed experimental evidence and best-fitted with computational models. The resultant online resource provides an effective reference to design new experiments, analyses, and spiking neural network simulations. Here we illustrate the content and utility of Hippocampome.org with a focus on the subiculum, whose neuron type organization has received relatively less attention. Only 6 of the 180 Hippocampome.org neuron types are from the subiculum, compared to more than 60 in the adjacent area CA1. Specifically, we analyze the local subicular circuit and its broader interaction with the hippocampal formation with respect to both anatomical connectivity and signal transfer. Our results exemplify the potential added value of data integration in neuronal classification, while also highlighting the need for further research to fill existing knowledge gaps.

## Introduction

The subiculum has been gaining recognition for its contribution to multiple cognitive tasks, like learning and memory (Böhm et al., 2018; Imbrosci et al., 2021) and goal-directed behavior (Lee & Lee, 2025), as well as its involvement in neurological disorders such as epilepsy (Baset & Huang, 2024; Knoppet al., 2005; Lévesque & Avoli, 2021; Lippmann et al., 2022). Anatomically, the subiculum constitutes a crucial nexus between area CA1 and the entorhinal cortex, thus playing a central communication role within the hippocampal formation (Aggleton & Christiansen, 2015). However, the neuron type organization of the subiculum remains understudied, especially relative to the surrounding regions of the hippocampus and neocortex (Jiang et al., 2025; Qiu et al., 2024).

Over the past 15 years, we have been developing Hippocampome.org, a web-accessible, open-source knowledge base of neuronal properties in the rodent hippocampal formation, including dentate gyrus, Cornu Ammonis, subiculum, and entorhinal cortex. Hippocampome.org categorizes glutamatergic and GABAergic neuron types primarily based on their axonal and dendritic patterns across distinct subregions and layers (Wheeler et al., 2015). This classification strategy is instrumental in identifying the mesoscale circuit organization in terms of potential connectivity (Ascoli & Wheeler, 2016; Rees et al., 2016) and helps resolve confusing terminological inconsistencies in naming neurons in the scientific literature (Hamilton et al., 2017a). This online resource further annotates each neuron type with rich multimodal properties, such as molecular expression (White et al., 2020), firing patterns (Komendantov et al., 2019), preferential spiking phases relative to in vivo rhythms (Sanchez-Aguilera et al., 2021), and cell counts (Attili et al., 2022). Moreover, directional interactions between neuron types are quantified in terms of synaptic physiology (Moradi & Ascoli, 2020) and connection probabilities (Tecuatl et al., 2021b). Each measurement reported in the database is directly linked to the corresponding published evidence, with full bibliographic reference and relevant metadata, tallying more than 520,000 pieces of knowledge and 46,000 pieces of evidence obtained from over 1500 peer-reviewed publications (Wheeler et al., 2024). Importantly, Hippocampome.org leverages these experimental data to constrain computational models of neuronal excitability (Venkadesh et al., 2019) and synaptic signaling (Moradi et al., 2022), thus providing a complete set of parameters to implement spiking neural network simulations (Sutton & Ascoli, 2021). The corresponding software tools are also freely released at the site, allowing users to customize the parameter fitting as needed. For example, the Synapse Modelers Workbench (Moradi et al., 2022) can adjust the amplitude, duration, and short-term plasticity of synaptic signals based on parameter settings such as recording modality (voltage vs. current clamp) and temperature, as well as the animal species (rat vs. mouse), age, and sex (Miller & Aoto, 2025). Similarly, the Connection Probabilities calculator (Tecuatl et al., 2021a) can update the connection probabilities between neuron type pairs if users provide custom estimates for dendritic spine density, inter bouton-distance, and interaction distances, which may depend on animal sex, age, pathologies, and more (Renner & Rasia-Filho, 2023). Humans and machines can mine this wealth of data respectively via graphical user interface or application programming interface (Nadella et al., 2025).

Out of 180 neuron types in Hippocampome.org, only 6 are from the subiculum, compared to over 60 in neighboring CA1. This might be due in part to the less apparent laminar architecture of the subiculum, which lacks the clear anatomical delineation of distinct input pathways in CA1, such as the CA3 Schaffer collaterals in stratum radiatum and the entorhinal perforant pathway in stratum lacunosum-moleculare (Gao et al., 2025). Yet, somatic expressions of parvalbumin, somatostatin, and vasoactive intestinal peptide (Kim et al., 2017), as well as broader transcriptomics evidence (Muñoz-Castañeda et al., 2024), suggest that the subiculum, like CA1, should also contain diverse populations of inhibitory interneurons as reported in all other hippocampal subregions. At a minimum, we would expect in the near future the fuller characterization in the subiculum of perisomatic-targeting, fast-spiking axo-axonic and basket cells (Okur & Scheiffele, 2025), input-specific dendritic targeting interneurons like CA1 O-LM, dentate HIPP, or cortical Martinotti cells (Park et al., 2025), and disinhibitory GABAergic-targeting interneurons (Guet-McCreight, et al., 2020). Evidently, the information currently available on subicular interneurons is still scarce.

Here we offer a brief overview of Hippocampome.org content focusing on properties of the neuron types identified in the subiculum so far. Additionally, we leverage available neurite reconstructions to quantitatively analyze neuron type circuit connectivity within the subiculum in terms of connection probabilities, number of contacts, and somatic path distances along the axonal and dendritic arbors. Moreover, harnessing data and tools from Hippocampome.org, we estimate the excitatory synaptic charge transfer between subicular neuron types and their input and output partners across the hippocampal formation, emphasizing key interactions with CA1 and entorhinal cortex. These applications illustrate the potential utility of neuroinformatics databases to foster further research into the subiculum neuron type circuit and explore its roles in important cognitive tasks.

## Material and Methods

### Hippocampome.org parameters

To analyze local circuit connectivity of subicular neuron types, we started from the connection probabilities and numbers of contacts reported at Hippocampome.org (RRID:SCR_009023). Similarly, we extracted synaptic physiology values for pairs of connected subicular neurons from hippocampome.org/php/synaptome_modeling.php selecting the following metadata: Species = rat; Sex = male; Age = P56; Temperature = 32 Celsius; Recording Mode = Voltage-clamp. We then augmented these initial datasets with additional measurements following the Hippocampome.org data analysis pipeline as described below. Readers are welcome to try variations of this analysis with alterative settings (e.g., P14 female mouse at 22 Celsius in current-clamp).

### Axonal and dendritic path distances

We used 1 to 3 two-dimensional reconstructed morphologies per identified subiculum neuron type and Simple Neurite Tracer (RRID:SCR_016566; Longair et al., 2011) to estimate average synaptic path distance from the soma along the axons and dendrites, as previously documented (Tecuatl et al., 2021a).

### Neurite length, convex hull, and potential connectivity

We previously reported the calculation of dendritic and axonal lengths from two-dimensional reconstructions (Tecuatl et al., 2021b). Here we apply this method to 10 three-dimensional reconstructions of subicular neurons from the MouseLight archive (Winnubst et al., 2019) of NeuroMorpho.Org (RRID:SCR_002145). Specifically, 4 files corresponded to CA1-projecting pyramidal-cells (AA0058, AA0820, AA0890, AA0894) and 6 to the distinct EC-projecting pyramidal neuron type (AA0024, AA0030, AA0032, AA0033, AA0703, AA0822), as mapped with the 2022 version of the Allen Institute Common Coordinate Framework (CCF). All values were corrected for the mouse-to-rat species conversion factor (Tecuatl et al., 2021b). We then extracted parcel-specific convex hull volumes from the same reconstructions using the procedure detailed previously (Tecuatl et al., 2021a). From these data, we then calculated the connection probabilities per directional pair of presynaptic and postsynaptic neuron types and the number of synaptic contacts in each anatomical layer as specified in the references cited above.

### Synaptic Charge transfer

We used the Hippocampome.org high-performance Synapse Modelers workbench (under the Tool menu) to estimate the charge transfer for each pair of connected neurons by exporting synaptic event traces from the available biophysical parameters in voltage clamp mode. Specifically, we first imported into the Synapse Modelers workbench a sample trace (github.com/k1moradi/SynapseModelingUtility - file name: 2100003). Next, we inserted in the graphical user interface the synaptic physiology parameters for each directed neuron type pair from Hippocampome.org (*g*_*syn*_,□_*d*_, □_*r*_, □_*f*_, *U*). We then selected the AMPA reversal potential (*E*_*rev*_□=□0□mV) and inserted the resting potential (*V*_*rest*_) of the postsynaptic neuron type from Hippocampome.org/php/ephys.php as the clamping voltage value (*V*_*m*_). With these inputs, the Synapse Modelers workbench generates a function of current over time as a comma-separated value file that we used to calculate the synaptic charge transfer *Q*_*s*_ as the area under the curve: *Q*_*s*_ = ∫ *I* (*t*) *dt* = ∫ *g* (*t*) * (*V*_*m*_ − *E*_*rev*_) *dt*

## Results

Hippocampome.org identifies 180 neuron types throughout the rodent hippocampal formation (Wheeler et al., 2024): 36 in the dentate gyrus (DG), 34 in CA3, 5 in CA2, 63 in CA1, 6 in subiculum, and 36 in the entorhinal cortex (EC). Each neuron type is annotated with properties, or features, across multiple dimensions, including morphological (e.g., subicular Wide-arbor Polymorphic interneurons have dendrites in the pyramidal layer…), molecular (… they express somatostatin…), electrophysiological (… have an input resistance of 387.8 ± 47.2 MΩ), and more. Users can browse neuronal properties based on a set of interactive matrices where neurons are listed as rows and properties as columns. Additional browsable matrices summarize synaptic properties where rows and columns represent presynaptic and postsynaptic neuron types, respectively. Each of over half a million properties are hyperlinked to peer-reviewed experimental evidence in the form of text excerpts, figures, and tables with detailed bibliographic references. The graphical user interface also enables searching neuron types based on arbitrary selections of properties.

Figure 1 illustrates a (very partial) view of this organization, as exemplified for the subiculum, starting from representative reconstructions of one of the glutamatergic neuron types, namely the EC-Projecting Pyramidal cells (Harris & Stewart, 2001). Notably, the two excitatory neuron types Hippocampome.org identifies in the subiculum have similar axonal and dendritic distribution within the subicular layers (Harris et al., 2001). However, they can be clearly distinguished by their stark axonal projection preferences for CA1 or EC, respectively. In contrast, GABAergic interneurons display more diversified local morphological patterns, as apparent when comparing the Pyramidale Multipolar cells, with axons and dendrites entirely contained within the stratum pyramidale (SP), against the Recurrent Polymorphic-Moleculare cells, with dendrites in the SP and the polymorphic layer (PL) and axons invading all 3 subicular layers. Although many more neuron types have been characterized in CA1 than in the subiculum, progress is still ongoing: the initial version 1.0 of Hippocampome.org (Wheeler et al., 2015) only recognized three subicular neuron types, with three new entries (marked with an asterisk in Fig. 1) added in the most recent release (Wheeler et al., 2024). On the transcriptomics front, both excitatory cell types display nearly identical molecular marker profile for calcium binding proteins, receptors, neuropeptides, and other biochemicals, whereas expression data on inhibitory interneuron types remains sparse. Based on similarity with CA1, we expect the future inclusion of distinct parvalbumin-positive and cholecystokinin-positive basket cells (Losonczy et al., 2002), Cajal-Retzius cells (Anstötz & Maccaferri, 2020), Ivy cells (Fuentealba et al., 2008), and others (Kawaguchi & Hama, 1987).

**Figure 1.**
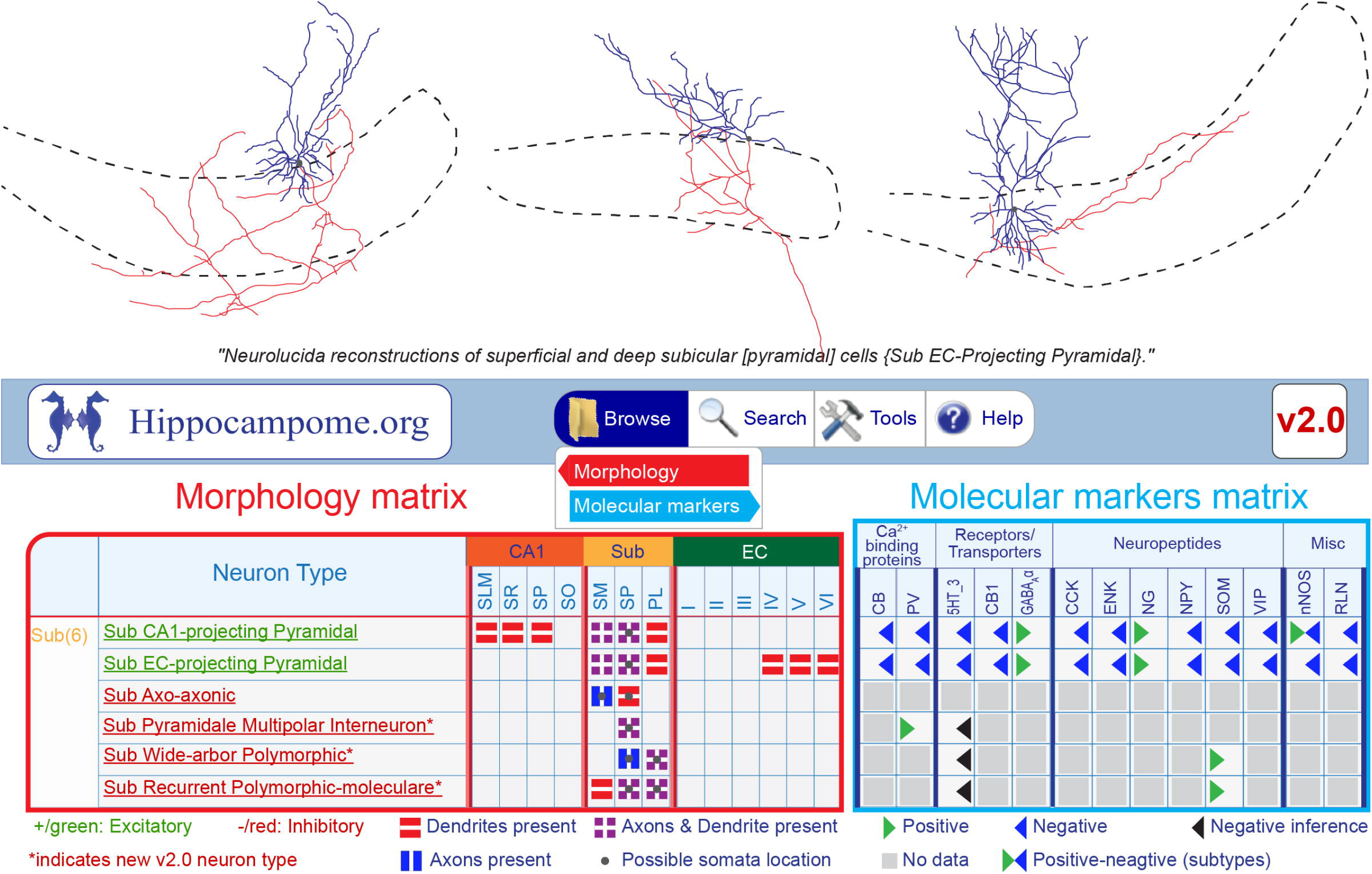
From morphology to neuron types. Top. Representative morphological reconstructions of subicular pyramidal neurons (Harris & Stewart, 2001) and related (quoted) caption, as included in Hippocampome.org, with somas in dark gray, dendrites in blue, axons in red, and dashed layer boundaries. The term in square parentheses is added for context from the cited reference, while the terms in curly brackets indicate the Hippocampome.org neuron name. Bottom. Hippocampome.org graphical user interface. Left. Portion of the morphology matrix (with red border) representing six subicular neuron types with laminar presence of axons (horizontal red stripes), dendrites (vertical blue stripes), both axons and dendrites (purple crossings), and somas (gray dot). Right. Portion of the molecular marker matrix (with blue border) for the same subicular neuron types. Abbreviations: CB, calbindin; PV, parvalbumin; 5HT_3, serotonin receptor type 3; CB1, cannabinoid receptor type 1; CCK, cholecystokinin; ENK, enkephalin; NG, neurogranin; NPY, neuropeptide Y; SOM, somatostatin; VIP, vasoactive intestinal peptide; nNOS neuronal nitric oxide synthase; RLN, reelin.

In addition to property-specific browse matrices, Hippocampome.org provides an individual page for each neuron type, summarizing all its features across dimensions, including axonal and dendritic distributions, molecular expression, membrane biophysics, firing patterns, cell counts, in vivo rhythms, input and output potential connectivity with other types, and more (Figure 2). These neuron type pages also list the neuron type name derivation, its synonyms, and supertypes, as well as bibliographic references, representative figures, and any curation note to facilitate interpretation (Hamilton et al., 2017a). As in the browse matrices, all information in these pages is hyperlinked to the underlying experimental evidence. Furthermore, the neuron type pages offer downloadable and ready-to-use electrophysiological simulation files for both the XPP (Anderson et al., 2015) and CarlSim (Niedermeier et al., 2022) simulators using Izhikevich models best fitted to the available experimental data (Venkadesh et al., 2018). Alternatively, Hippocampome.org allows users to launch these models directly in their web browser, adjust parameters as desired, and plot or save the results on their local devices. The neuron type pages further fetch relevant digital reconstructions of neuronal morphology from NeuroMorpho.Org (Tecuatl et al., 2024) and compartmental Hodgkin-Huxley models from ModelDB (McDougal et al., 2017). This approach extends the dense coverage of current knowledge from the peer reviewed literature by maximizing the reusability of data from complementary open access resources. As an example, Figure 2 illustrates a portion of the subiculum CA1-Projecting Pyramidal cell Hippocampome.org page, which gleans 133 properties mined from 31 publications.

**Figure 2.**
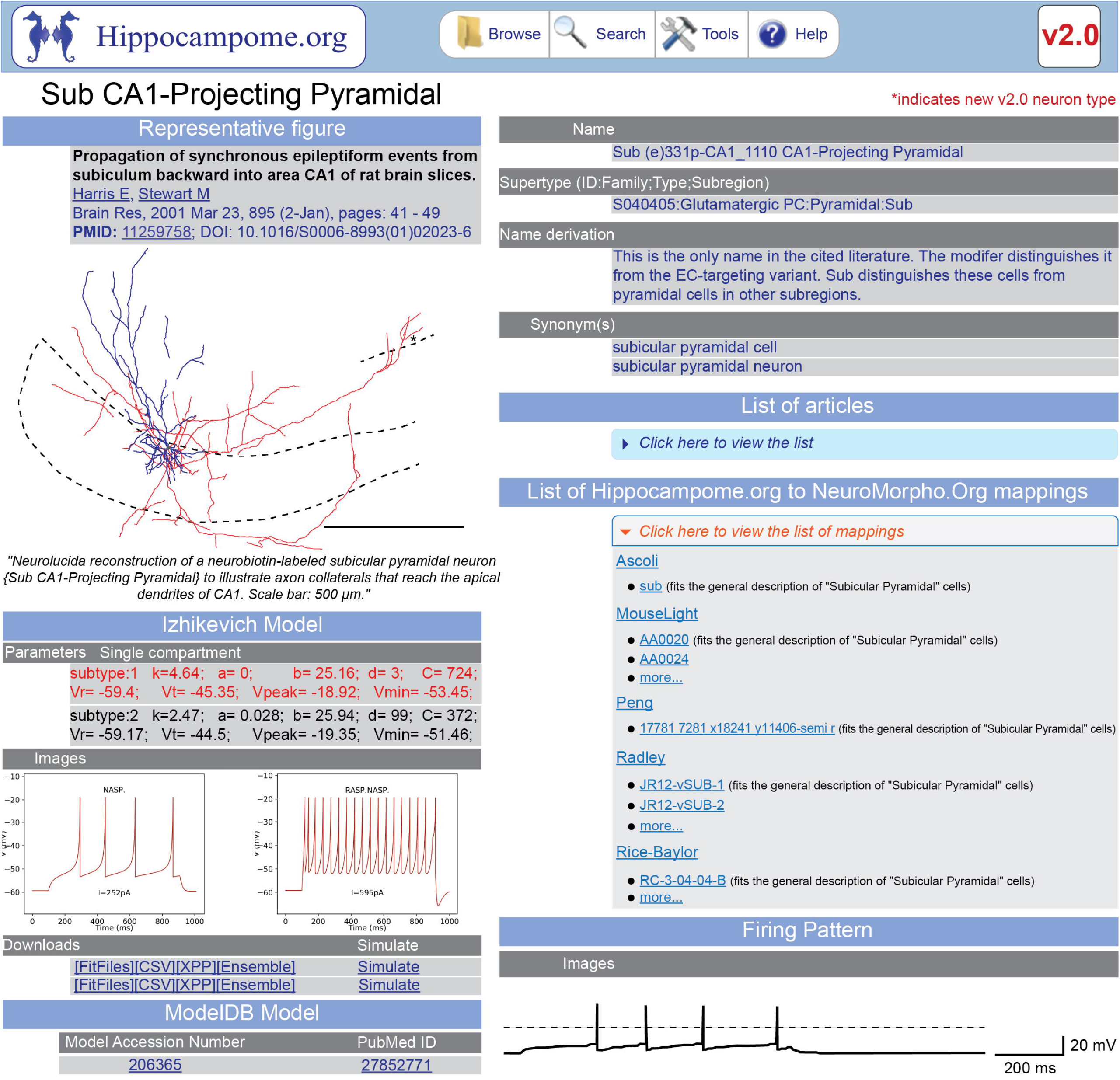
Excerpt of the Hippocampome.org subiculum CA1-Projecting Pyramidal neuron type page. Left. Representative figure with linked reference (original colors were modified to show axons in red and dendrites in blue) and quoted caption; below are Izhikevich model parameters for two electrophysiological subtypes of this neuron type with corresponding simulated firing patterns, downloadable and executable simulation files, and linked ModelDB entries. Right. General information (name and derivation notes, supertype, and synonyms), clickable list of articles describing that neuron type (not expanded in this illustration), hyperlinked mapping to NeuroMorpho.Org reconstructions, and experimentally recorded firing pattern.

Next, we turn our attention to circuitry, capturing the directional interaction between neuron types. The laminar distributions of axons and dendrites (and the postsynaptic somatic location for perisomatic targeting interneurons) are sufficient to determine which pairs of pre- and post-synaptic neuron types can form potential connections. Starting from this foundation, Hippocampome.org quantifies multiple connectivity features, including the average synaptic distances from the soma along the axonal and dendritic path, the connection probability, and the mean number of contacts per connected neuron pair (Figure 3). Previously, we only measured these values within local circuits due to major axonal projection loss in typical slice preparations. Moreover, Hippocampome.org only provided quantitative connectivity data for the three subicular neuron types included in Hippocampome.org v.1.0, given the relative sparsity of earlier published data. Here, we add the measurements for the other three subicular neuron types available in Hippocampome.org v.2.0, yielding 11 new somatic distances obtained from 4 two-dimensional reconstructions. Figure 3A displays representative results for the EC-Projecting Pyramidal neuron, while all others are provided in the Supplementary information as detailed in the Data Sharing and Data Accessibility section below. The added morphological tracings also allow a substantial expansion of the quantitative estimation of the local subicular circuit connectivity, because two out of the three new interneuron types extend dendrites in the polymorphic layer, an anatomical parcel which previously lacked identified dendritic targets (Fig. 3B). The connection probabilities vary widely across the 25 neuron type pairs (mean ± standard deviation: 0.4% ± 0.1%, median 0.1%) with a minimum of 0.0065% from subiculum Pyramidale Multipolar interneuron to the CA1-Projecting Pyramidal cells and a maximum of 2.5% between CA1-Projecting Pyramidal cells. The number of contacts per connected pair in the local subiculum circuit, 6.46 ± 1.46, falls within the range of values observed in other hippocampal regions (Tecuatl et al., 2021b). Breaking down the number of contacts by presynaptic and postsynaptic types yields 4.5 ± 1.59 for excitatory-excitatory pairs (n=4), 2.7 ± 0.53 for excitatory-inhibitory (n=8); 11.1 ± 3.23 for inhibitory-excitatory (n=4), and 8.6 ± 4.21 for inhibitory-inhibitory (n=9), confirming previous findings of larger values for inhibitory presynaptic pairs (Tecuatl et al., 2021b).

**Figure 3.**
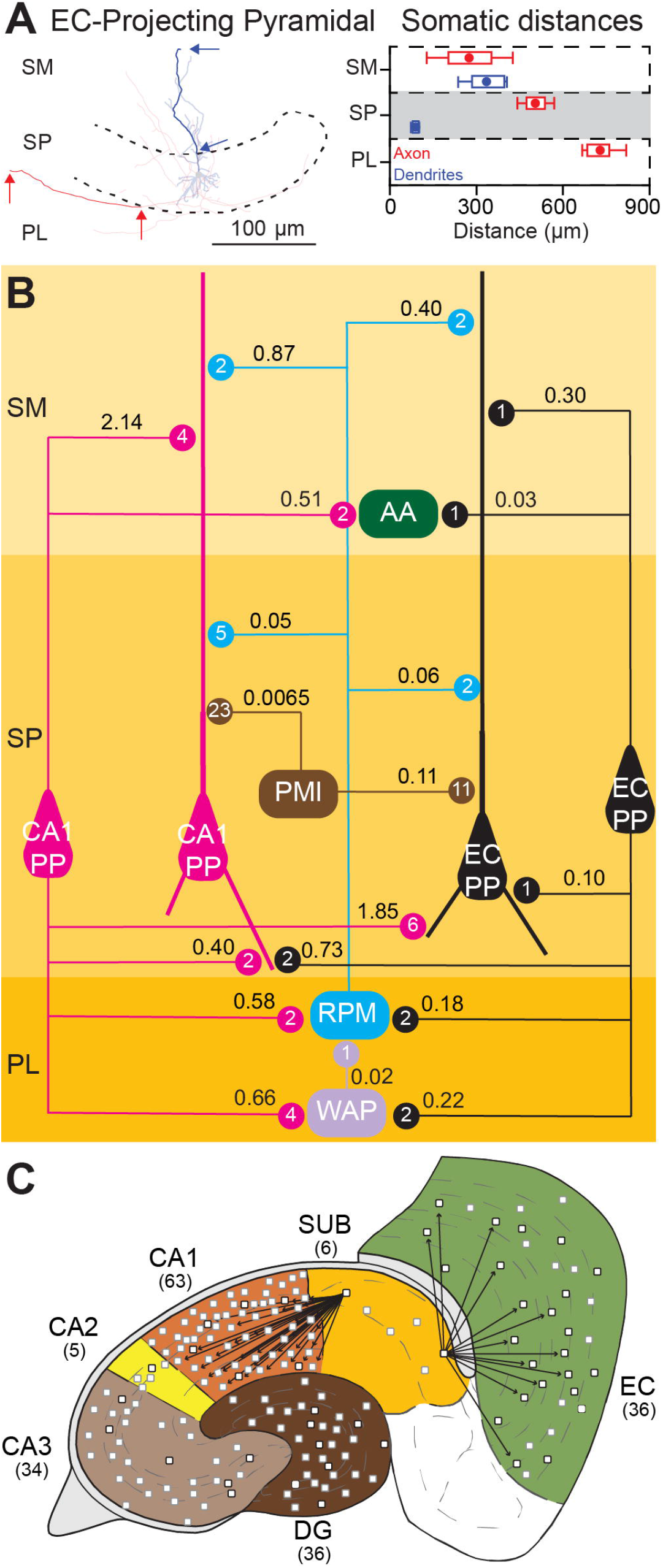
Subicular connectivity. A. Left. Schematics of dendritic and axonal path distances from the soma across different subiculum layers from a morphological reconstruction of EC-Projecting Pyramidal cell. Right. Average somatic distances along axonal (red) and dendritic (blue) paths by subicular strata obtained from two EC-Projecting Pyramidal reconstructions (n=4 dendritic paths, n=2 axonal paths). B. Quantification of the local subiculum neuron type circuit (thin lines: axons; thick lines: dendrites). Connection probabilities are indicated along the axonal lines, while the numbers of contacts per connected neuron pair are shown in circles. Abbreviations: CA1, Cornu Ammonis area 1; CA2, Cornu Ammonis area 2; CA3, Cornu Ammonis area 3; EC, entorhinal cortex; CA1 PP, CA1-Projecting Pyramidal; EC PP, EC-Projecting Pyramidal; PMI, Pyramidale Multipolar interneuron*; RPM, Recurrent Polymorphic-Moleculare*; WAP, Wide-arbor Polymorphic*; PL, polymorphic layer; SP, stratum pyramidale; SM, stratum moleculare. C. Schematic representation of subicular projections to 31 CA1 and 13 EC neuron types.

Furthermore, it has recently become possible to reconstruct projection neurons with full axonal extent mapped to a common coordinate framework (CCF). These technological advancements make it feasible to estimate probabilities of connection from major projection neuron types. Here, we quantify the projection connectivity from the two (CA1-projecting and EC-projecting, respectively) subiculum pyramidal cells (Fig. 3C). For this purpose, we downloaded from the MouseLight archive of NeuroMorpho.Org (Winnubst et al., 2019) 10 reconstructions with neurite patterns corresponding to those reported by Hippocampome.org when mapped to the CCF v2.0 anatomical parcels (Wang et al., 2020). Species-normalized, parcel-specific axonal lengths are reported in Supplementary Table 1. With these new measurements, we refined 29 connection probabilities in CA1 previously estimated from 2-dimensional reconstructions that were confined to strata lacunosum-moleculare (SLM) and radiatum (SR) but missed SP, and added a new connection probability. Furthermore, we added 13 overall and 23 parcel-specific connection probabilities in EC. Results for the connectivity from the subiculum CA1-Projecting Pyramidal cell to 31 distinct CA1 neuron types show a percentual connection probability of 0.20 ± 0.03 with 4.6 ± 0.46 contacts per connected pair. The parcel specific analysis reveals the highest percentual probability in SP (0.20 ± 0.015; n=15) and the lowest in SLM (0.04 ± 0.005; n=20), with an exactly opposite relation for the number of contacts (3.5 ± 0.55 in SLM vs. 1.0 ± 0.06 in SP). This latter trend is consistent with previous findings of more dendritic spines in SLM than in SP in CA1 pyramidal cells (Megí as et al., 2001). Interestingly, the subiculum EC-Projecting Pyramidal cells display a remarkably higher percentual connection probability to the 31 target neuron types in EC (12.0 ± 2.93), with 2.2 ± 0.33 contacts per connected pair. These data contribute much needed information to create spiking neural network models of the subiculum. However, several connection probabilities from the subiculum to EC are still missing due to the lack of sufficiently complete dendritic reconstructions for MEC LIII Multipolar Principal cells, MEC LIII Multipolar interneurons, and EC LIII Pyramidal-Looking interneurons. Similarly, the dearth of fully reconstructed axonal projections from EC to the subiculum prevents their connectivity quantification.

Finally, we turn our attention to the analysis of excitatory synaptic transmission between specified pre-synaptic and post-synaptic neuron types, while maintaining the focus on the subiculum (Figure 4). In particular, we quantify the synaptic charge transfer, or total amount of electrical signal recorded during a synaptic event (Turner & Schwartzkroin, 1983), computed as the integral of current over time (Fig. 4A). Figure 4B graphs the synaptic charge transfer network of 72 neuron type pairs, corresponding to 6 connections within the local subicular circuit, 27 afferents to the subiculum (21 from EC, 3 from CA3, and 3 from CA1), and 39 efferents from the subiculum (22 to CA1, 16 to EC, and 1 to CA2). Here we aggregate the values as average ± standard error over the number of synaptic pairs from selected subcircuits. Overall, the synaptic charge transfer for neuron type pairs within the local subiculum circuit (-1788.26 ± 104.7 pC, n=6) is statistically similar to that for afferent inputs to the subiculum (-1750.54 ± 37.5 pC, n=27: p>0.5). In contrast, the efferent outputs from subiculum have significantly lower synaptic charge transfer (-1584.61 ± 41.6 pC, n=39: p<0.05). Looking into more details, the synaptic charge transfer from CA1 to the subiculum (-1595.6 ± 108.3 pC, n=3) is statistically similar to that from the subiculum to CA1 (-1669.8 ± 65.9 pC, n=22; p>0.5). This seems consistent with the view of the subiculum playing a tight regulatory role on local CA1 activity (Gao et al., 2025). The relationship between the EC and the subiculum is remarkably different. The synaptic charge transfer from the EC to the subiculum (-1772.0 ± 42.5 pC, n=21) is significantly higher than from the subiculum to the EC (-1468.4 ± 28.2 pC, n=16, p<0.001), which further challenges the canonical view of directional information flow in the hippocampal formation (Farrell & Soltesz, 2025). Last but not least, we observe that the excitatory synaptic charge transfer on postsynaptic GABAergic neurons (-1759.0 ± 53.6 pC, n=33) is significantly larger than on postsynaptic glutamatergic neurons (-1583.2 ± 22.2 pC, n=39; p<0.005), a trend that holds for all subcircuits analyzed in this study. This difference may be related to preferential subunit expression of ionotropic glutamate receptors in distinct neuron types (Hamilton et al., 2017b; Yao et al., 2021).

**Figure 4.**
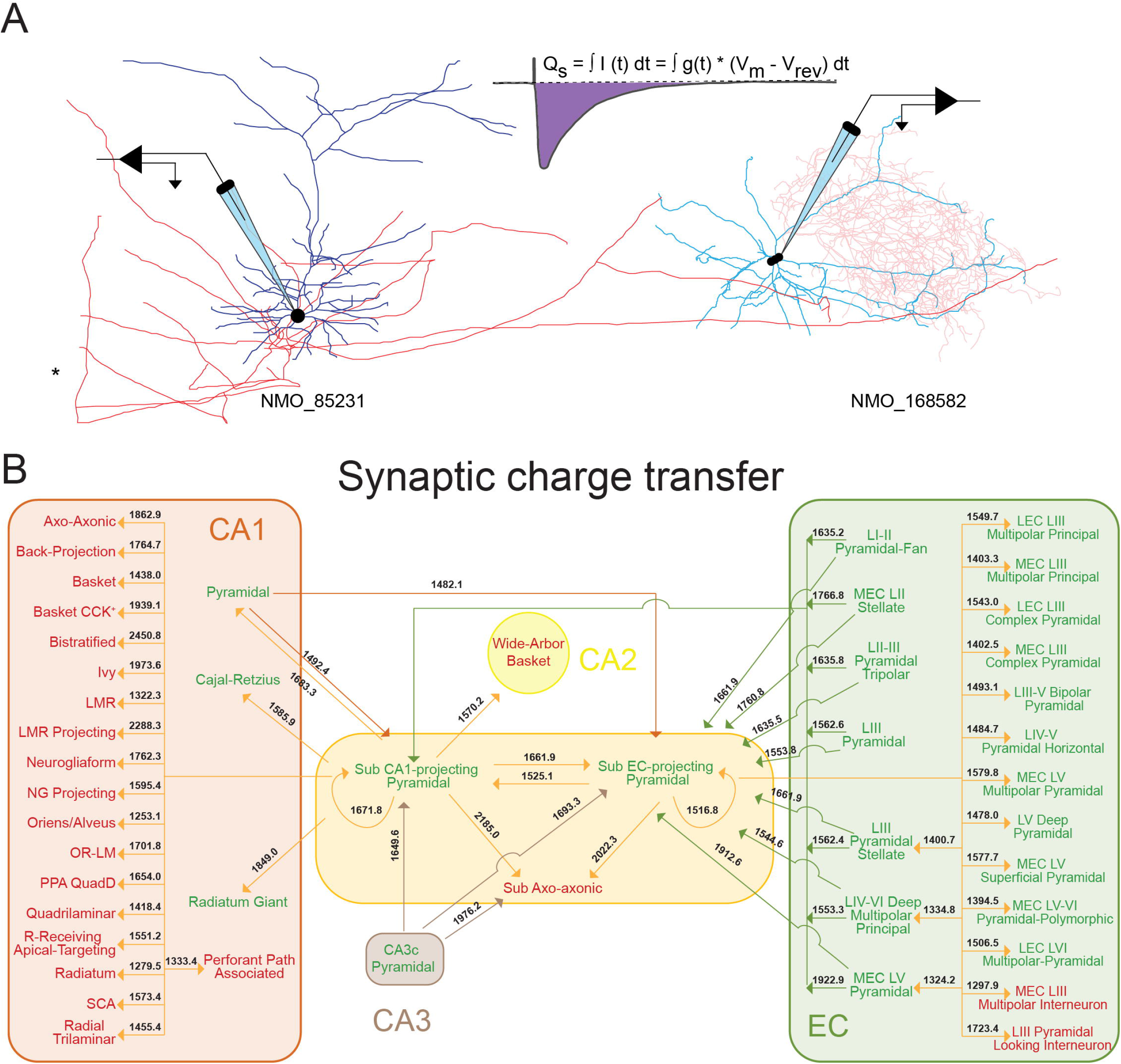
Subicular synaptic charge transfer. A. Representative pair of synaptically connected subicular neurons: a pyramidal cell (Gerfen et al., 2018), left, contacting an axo-axonic interneuron (Anstötz et al., 2021), right. Reconstructions were obtained from NeuroMorpho.Org, after removing the long-range projection portion of the pyramidal cell axons for clearer illustration of the local subiculum circuit (red: axons; blue: dendrites; magenta: apical dendrite). B.Connectivity diagram of the subicular excitatory synaptic charge transfer network. Rounded boxes represent the indicated regions of the hippocampal formation centered around the subiculum (yellow). Arrows are colored by presynaptic region, with arrowheads pointing in the synaptic flow direction and the numbers on the lines indicating the absolute values of the synaptic charge transfer in pC. Glutamatergic and GABAergic neuron types are shown in green and red, respectively. Abbreviations: CA1, Cornu Ammonis area 1; CA2, Cornu Ammonis area 2; CA3, Cornu Ammonis area 3; EC, entorhinal cortex; MEC, medial entorhinal cortex; LEC; lateral entorhinal cortex; NG, neurogliaform; PPA, perforant path-associated; SCA, Schaffer collateral-associated.

Altogether, these results contribute to Hippocampome.org 600 new pieces of knowledge and 459 pieces of evidence, all available as supplementary data. The correspondingly expanded characterization of subicular neuron types and of their circuitry constitutes a substantial steppingstone towards a fuller understanding of their functional roles. At the same time, it is clear that the expected discovery and characterization of additional neuron types in the subiculum will further enrich this evolving knowledge base. Likewise, synaptic information is still missing for the connections from at least 8 CA1 neuron types and 7 EC neuron types to the subiculum according to the Hippocampome.org potential connectome. Thus, new pieces of information will be required to achieve a relatively complete computational model of the subicular circuit, including representative morphological reconstructions, electrophysiological recordings, neuron type-specific population counts, and more.

## Discussion

Neuroscientists no longer overlook the subiculum as a mere output of the hippocampal formation to neocortical and subcortical areas, as demonstrated by the increasing number of publications that reveal subicular roles in multiple cognitive processes (Place et al., 2025). Recent discoveries include active boundary vector cells in the dorsal subiculum that align with future (rather than current or past) positions during spatial navigation (Newman et al., 2025; Poulter et al., 2021), and the aging-related decline in the neural representation of boundary information in both EC and subiculum (Segen et al., 2025). Nevertheless, this rising scientific interest in subicular function has yet to stimulate more intense research on the characterization of neuron types, their properties, and circuit connectivity. Our latest analysis of web traffic at Hippocampome.org still shows that public interest in subiculum neuron types is not even close to that in more studied brain regions like CA1 or DG (Nadella et al., 2025). Thus, a widening gap appears to exist between accumulating knowledge on the multifaceted contributions of the subiculum to complex behaviors and the scarcity of detailed data on the underlying neuronal organization.

This article highlighted the potential utility of Hippocampome.org in studying neuron type architecture, focusing on the subiculum as the driving example. With sufficient information, this resource may provide all data-driven parameters necessary to create real scale, biologically realistically spiking neural networks, as recently demonstrated for area CA3 (Kopsick et al., 2023). Those computational simulations are useful tools to test and refine theories relating specific circuit activity patterns with cognitive functions, such as cell assemblies in pattern completion and associative retrieval (Kopsick et al., 2024), and continuous attractor dynamics in trajectory coding (Rg et al., 2025; Sutton et al., 2024). Reaching a similar state of maturity for the subiculum, however, will require the discovery of new subicular neuron types and more comprehensive characterization of existing ones.

For instance, although subiculum CA1-Projecting Pyramidal cells can stimulate basal dendrites in SO (Köhler, 1985), the extent of subicular axonal morphology actually reconstructed to date only includes CA1 projections to SLM, SR, and SP. Similarly, subiculum EC-Projecting Pyramidal cells contact dendritic targets in EC LIII (Kloosterman et al., 2000; Kloosterman et al., 2004; Kloosterman et al., 2003), but existing axonal reconstructions are limited to layers IV-VI. More generally, the Hippocampome.org classification of subiculum long-range pyramidal cells assumes that each neuron preferentially targets a single region (CA1 or EC) without noting possible laminar selectivity (Witter & Groenewegen, 1990). We need more data to improve this likely over-simplified view in light of early evidence on subicular neurons selectively targeting CA1 SO or with axons collateralizing in both CA1 and EC (Finch et al., 1983). More recent reports describe other subicular neurons with axons reaching CA2, CA3, and DG (Winnubst et al., 2019) or with preferential projections to either LEC or MEC (Qiu et al., 2024).

The above observations support the existence of new cell types, possibly related to distinct transcriptomics profiles (Ding et al., 2020). Nonetheless, when considering long-distance axonal tracings, larger sample sizes and inter-laboratory reproducibility remain essential to carefully rule out the ever-present risk of misaligned anatomical mapping due to differences in individual brain shape, age, and other processing factors like tissue shrinkage and histological damage (Qiu et al., 2024). A recent analysis of an adjacent region, the presubiculum, provides a compelling demonstration of the power of massive, brain-wide neuronal morphology datasets to understand circuit logic (Wheeler et al., 2024). In that case, the large number of nearly complete axonal reconstructions from the presubiculum enabled the deployment of a data-driven, unsupervised clustering of projection patterns (Wheeler & Ascoli, 2025) that revealed five different neuron types with highly specific anatomical targets.

The discovery and characterization of new GABAergic interneuron types is also necessary to better dissect the local subicular circuit. Transgenic mice have proven useful to investigate specific neuron populations and could provide a productive approach to identify similarities and differences with cell types in other hippocampal regions (Anstötz et al., 2021). In terms of functional synaptic connectivity, the data for subicular neuron types is even more scant, and largely derived from indirect evidence. For example, recordings from postsynaptic subiculum targets upon stimulation of CA1 SR (Price et al., 2005) or SLM (Milstein et al., 2015) fibers could be ascribed to any presynaptic neuron with axons in those areas. Similarly, we could not map an elegant study of excitatory/inhibitory interplay in subicular neurons (Böhm et al., 2015) to identified pre-/post-synaptic neuron types given the lack of specific GABAergic markers and the similar morphological, molecular, and electrophysiological properties of EC- and CA1-projecting pyramidal cells (Harris & Stewart, 2001; Kim & Spruston, 2012).

In this work, we attempted to expand the nascent characterization of the subiculum neuron type circuit by filling selected gaps for 72 pairs of potentially connected neuron types through a neuroinformatics strategy. In particular, we leveraged available data from NeuroMorpho.Org along open-source toolset and existing content of Hippocampome.org to quantitatively estimate both structural (connection probability and number of contacts) and functional (synaptic charge transfer) parameters. Although this is a relatively small step in the long path to collecting all missing information, it is still a substantial contribution that would have been difficult and costly to achieve with a direct experimental approach. We hope these results may aid future analysis and modeling efforts in shedding light on the neuron circuit basis of subicular function.

In conclusion, Hippocampome.org constitutes a rich knowledge base for the study of neuron types, their properties, and synaptic communication throughout the rodent hippocampal formation. The additional data generated for the subiculum through the latest Hippocampome.org release and in the work presented here considerably increased previously available information. However, a remarkable gap still exists between the subiculum and other hippocampal regions in the amount of published neuron type data. As evidence continues to accumulate on the subicular involvement in many cognitive and behavioral tasks, it is urgent to advance towards the discovery and characterization of additional neuron types in this brain region. This will enable the implementation of real-scale spiking neural network models that may help anchor computational functions with neuronal, synaptic, and circuit mechanisms.

## Supporting information

Connectivity summary

Synaptic charge summary

## Acknowledgments

The authors are grateful to Keivan Moradi, Jeffrey D. Kopsick, and Diek W. Wheeler for their help with and feedback on data acquisition; and to Kasturi Nadella, Lin Shen, and Zengxin Li for support and maintenance of the computing resources.

## Data Sharing and Data Accessibility

All results and underlying experimental data are freely available through the Supplementary information. Specifically, the Connectivity folder contains a Connectivity_Summary Excel file that reports, for all pairs of potentially connected neuron types, the means, minima, and maxima of dendritic and axonal lengths; means, SDs of the synaptic distances from the soma along the axonal and dendritic paths; and means for the connection probabilities, average total numbers of synapses per neuron pair, and numbers of contacts per connected pair. Each set of values is linked to the underlying data in the Evidence folder, namely the morphological reconstruction in the Reconstruction subfolder and the convex hull images and path traces in the Convex_Hull subfolder. Synaptic charge transfer data are provided in their namesake folder that contains an Excel file with the synaptic modeler parameters. Model data for each synaptic pair are also provided in individual comma-separated-value (CSV) files in the corresponding Evidence folder. This material will also be accessible from hippocampome.org/php/help.php after manuscript publication.

## Author contributions

CT, GAA, Conception and design, Acquisition of data, Analysis and interpretation of data, Drafting, and revising the article.

