## Supplementary Table 1 for "Hippocampome.org, a resource for subicular neuron types and beyond"

Supplementary Table 1. Summary of total projection corrected axonal length of digital reconstructions obtained from NeuroMorpho.Org (MouseLight archive).

| NeuroMorpho.Org_ID | MouseLight_ID | Projection pattern | Axonal length (µm) | DOI |
| --- | --- | --- | --- | --- |
| NMO_85243 | AA0058 | CA1 | 4358.97 | 10.25378/janelia.7649933 |
| NMO_121856 | AA0820 |  | 4994.00 | 10.25378/janelia.7739729 |
| NMO_121862 | AA0890 |  | 6997.78 | 10.25378/janelia.7780769 |
| NMO_121820 | AA0894 |  | 6222.59 | 10.25378/janelia.7780793 |
| NMO_85233 | AA0032 | EC | 26609.96 | 10.25378/janelia.5521678 |
| NMO_121853 | AA0703 |  | 21271.65 | 10.25378/janelia.7704362 |
| NMO_85230 | AA0024 | MEC | 11736.03 | 10.25378/janelia.5521654 |
| NMO_85232 | AA0030 |  | 25115.28 | 10.25378/janelia.5521672 |
| NMO_85240 | AA0033 |  | 18848.46 | 10.25378/janelia.5521681 |
| NMO_121849 | AA0822 |  | 10743.53 | 10.25378/janelia.7739735 |
| NMO_85230 | AA0024 | LEC | 2461.91 | 10.25378/janelia.5521654 |
| NMO_85232 | AA0030 |  | 12279.42 | 10.25378/janelia.5521672 |
| NMO_121849 | AA0822 |  | 8550.46 | 10.25378/janelia.7739735 |
